# Wild chimpanzees in Bugoma Forest, Uganda follow the Ugandan feeding ecology etiquette but exhibit cultural singularities: a case for the cultural junction hypothesis

**DOI:** 10.64898/2026.03.10.710845

**Authors:** Kelly Ray Mannion, Catherine Hobaiter, Thibaud Gruber

**Affiliations:** Swiss Center for Affective Sciences, University of Geneva, Switzerland; Bugoma Primate Conservation Project, Kikuube District, Uganda; School of Psychology and Neurosciences, University of St Andrews, UK; Faculty of Psychology and Educational Sciences, University of Geneva, Switzerland

**Keywords:** ecology, chimpanzee, *Pan troglodytes schweinfurthii*, culture, biogeography

## Abstract

Chimpanzees, amongst other primates, are characterized by the large variability of habitats they can be found in as well as a large behavioral, sometimes cultural diversity. Such observations have launched a decades-long debate on the roots of behavioral diversity, stressing the need to document this behavioral variability in context, such as by connecting closely related populations through localized analyses. This study presents the first comprehensive description of feeding ecology from the Mwera South chimpanzee (*Pan troglodytes schweinfurthii*) community in the Bugoma Central Forest Reserve, in Uganda, establishing a valuable baseline for this previously unstudied population and providing a comparative perspective on the populations of Western Uganda and Eastern Democratic Republic of Congo. By employing multiple methodological approaches, including direct observation and fecal analysis, we describe dietary composition, seasonal patterns, and environmental influences on feeding behavior. Characterizing the feeding ecology of this previously unstudied population is critical for examining how ecological factors might influence how feeding patterns evolve depending on resource availability or chimpanzee behavior, in particular by favoring analyses at the regional level. In addition, we can better evaluate to what extent behavioral differences between chimpanzee communities stem from ecological constraints and/or cultural transmission pathways. Our findings suggest that the Bugoma chimpanzees seat at the location of a historic cultural junction, opening a large array of questions about historic movements and cultural diffusion in Eastern chimpanzees.

## Introduction

Understanding primate feeding ecology is fundamental to conservation biology and behavioral ecology, as dietary patterns influence individual survival, population dynamics, social behavior, and cultural transmission (Boesch et al., 2006; Wrangham et al., 1991). Primates, in particular great apes, exhibit complex feeding strategies that reflect both ecological constraints and behavioral, sometimes cultural, adaptations to their environment (Grund et al., 2019; Schuppli & van Schaik, 2019). Chimpanzees (*Pan troglodytes*) provide an excellent model for investigating the interplay between environmental conditions and feeding ecology, as they occupy diverse habitats across Africa and demonstrate considerable behavioral flexibility (Abwe et al., 2020; Ebang Ella et al., 2023; Kalan et al., 2020; Watts et al., 2012). However, despite extensive research at established sites, critical gaps remain in our understanding of how local ecological conditions influence feeding strategies and how these patterns relate to the behavioral diversity observed across populations.

This study presents the first comprehensive description of feeding ecology from the Mwera South chimpanzee (*Pan troglodytes schweinfurthii*) community in the Bugoma Central Forest Reserve, in Uganda. Doing so allows us to establish valuable ecological baselines for this previously unstudied population, and to provide a comparative perspective on the other well-studied communities in Uganda (Gruber, 2013, 2019). In addition, we address three critical aspects of primate diet, with each illustrating possible challenges facing researchers when evaluating, quantifying, and considering diet within broader contexts: i) environmental impact on diet and timescales; ii) methodological challenges, particularly when estimating diet in semi-habituated communities where gaps in data collection are frequent; and iii) cultural aspects of diet.

### Environmental aspects and timescales

Environmental factors significantly influence primate feeding patterns: both immediate weather conditions and broader seasonal changes affect food availability and feeding strategies (Grund et al., 2019; Potts et al., 2020). For example, seasonal variation in rainfall patterns affects fruit availability, while temperature and humidity can influence both food distribution and animal activity patterns (Potts et al., 2011). Chimpanzees typically respond to such variation through dietary switching, alternating between preferred foods (primarily ripe fruits) when available and fallback foods during periods of scarcity (Harrison C Marshall, 2011). However, the extent and nature of these dietary shifts can vary considerably between populations, reflecting both local ecological conditions and, potentially, cultural differences in feeding behavior (Gruber et al., 2012). Variation in feeding ecology and other intrinsic factors, such as travel effort, influenced interest in, engagement with, and use of tools in experimental set-ups that provide desirable foods, such as honey (Gruber et al., 2016; Grund et al., 2019). These effects occur over both the short-term (i.e. 2 to 7 days) and long-term timescales (i.e. 30 to 90 days), although not every effect is apparent at each timescale. For instance, in the Sonso chimpanzees of the Budongo Forest, engagement with the experiment was influenced in the long-term by a triple interaction between diet, travel effort, and duration (up to 90 days), but only travel effort predicted tool use at a shorter timescale (7 days; Gruber, et al., 2016). To address the influence of timescales, here we studied the effect of various environmental factors (temperature, humidity, rainfall etc.) on diet composition, particularly fruit feeding, using rolling averages at various timescales (from 2 to 90 days) to compare temporal variation, identify best fits, and study the wider trends that influence diet composition in the Mwera South chimpanzees of Bugoma Forest.

### Methodological challenges

Studies of primate feeding ecology increasingly recognize of the importance of multiple data collection methods (Rothman et al., 2012) to capture the full spectrum of feeding behavior. Observational techniques (Altmann, 1974) provide detailed behavioral data but may miss certain aspects of diet composition, particularly during periods when animals are difficult to observe. Complementary methods such as fecal analysis (McGrew et al., 2009) and eDNA analyses (Schneider et al., 2023) can provide additional insight into diet composition when studying unhabituated communities, or when logistical constraints prevent continuous observation (Basabose, 2002; Doran et al., 2002). They can help to validate observational data (Phillips & McGrew, 2013) as well as to identify food items consumed outside observation periods (Matthews et al., 2020; McLennan, 2013). However, comparing feeding ecology across sites presents particular challenges where methods or the combinations of methods differ (Potts et al., 2011), as well as where limited infrastructure prevents molecular-level or similarly detailed analyses. While we were unable to conduct eDNA analyses in this study, we compared records of direct observations of feeding behavior with fecal analyses. In particular, we investigated whether the consumption of specific items such as meat was correctly reflected in observational sampling.

### Cultural aspects

The Kibale National Park, home to several habituated communities, has revealed distinct dietary patterns despite the communities’ geographical proximity (Potts et al., 2011). For example, the Kanyawara chimpanzees show greater reliance on terrestrial herbaceous vegetation (THV) during periods of fruit scarcity, while the Ngogo chimpanzees maintain a more fruit-dominated diet throughout the year, likely due to higher fruit tree density in their home range (Watts et al., 2012). In contrast, the chimpanzees of the Budongo Forest Reserve experience different ecological challenges, with a more diverse forest composition and greater seasonal variation in food availability (Gruber et al., 2012). Here, the neighboring Sonso and Waibira communities show distinct differences in prey species consumption that are unexplained by ecological factors to date, suggesting cultural influences on feeding traditions (Hobaiter et al., 2014).

Bugoma Forest occupies a critical geographical position between these well-studied sites. Observations and field experiments suggest that Bugoma chimpanzees exhibit a mix of behavior observed in both Budongo and Kibale communities, including diverse tool use repertoires (Mannion et al., 2025) and ground nesting behavior more typically associated with Congolese communities to the west (Hobaiter et al., 2024; Romani et al., 2023), sitting at a potential “cultural junction” at which different behavioral traditions converge (Gruber, 2013; McLennan et al., 2020). Testing whether the feeding ecology of the Bugoma chimpanzees aligns with the one of the two neighboring populations (and if so, which) represents a first step, and will allow us to explore where and when ecological differences impact behavioral, and perhaps cultural, variation (Berger et al., 2019; Laland & Janik, 2006).

In this study we tested several key hypotheses about chimpanzee feeding ecology in the Bugoma Forest. With respect to environmental aspects, we tested (H1) that environmental conditions, particularly rainfall patterns, correlated with dietary choices; and (H2) that dietary diversity varied inversely with fruit availability, with higher dietary diversity during periods when fruit consumption is low. With respect to methodological challenges, we tested (H3) that fecal analysis and observational sampling provided complementary dietary data with methodological differences in detection capabilities, particularly for a community that was semi-habituated at the time of data collection. Finally, with respect to the cultural junction hypothesis, we tested (H4) that chimpanzee diets in the Bugoma Forest showed distinct compositional patterns characterized by strong species preferences, particularly figs, as found in other Uganda populations; and, more regionally, (H5) that the Mwera South community’s feeding patterns showed similarities with both Budongo and Kibale chimpanzee communities, in line with the cultural junction hypothesis.

## Methods

### Study Site

Bugoma Central Forest Reserve (CFR), Uganda (01°15′N 30°58′E) is a semi-deciduous tropical rainforest situated between Budongo Central Forest Reserve (425 km²) and Kibale National Park (776 km²). Since 2016, the Bugoma Primate Conservation Project (BPCP, www.bugomaprimates.com) has conducted systematic habituation and data collection from the Mwera field station (01°17’N 30°59’E) on the Mwera South community, which comprises approximately 70 individuals—a typical size in eastern chimpanzee (*Pan troglodytes schweinfurthii*) communities (Wilson et al., 2014), as well as with several other communities within the Bugoma CFR for shorter periods of time.

## Data Collection

### Diet data

Dietary data for the Mwera South chimpanzee community in the Bugoma Forest were collected through direct observations during habituation follows from April 2021 to June 2024. Field assistants and researchers systematically recorded the species and specific plant parts ingested during follows as daily presence/absence in the diet. Researchers and assistants typically left camp at 6am to find the chimpanzees and started to record data when the chimpanzees were located. Data collection continued throughout the day, while the chimpanzees were in sight, until approximately 5-6pm when the team returned to camp. Data collected throughout the study started off sparingly as it co-occurred with the progressive re-opening of Uganda to research following the Covid19 outbreak and the resumption of research activities. Time spent with the chimpanzees increased across the four years of data collection from 2021-2024, as research efforts could be increased and contact time and habituation improved. Overall, this represented 2373 feeding item records across 964 unique observation days.

### Weather data

We recorded daily weather measurements including Temperature (High, Low) and Humidity (High, Low) with a digital meter, and daily rainfall using a rain gauge. We used these parameters to analyze relationships between environmental conditions and feeding behavior.

### Fecal sample collections

We collected fresh fecal samples (N=75) opportunistically during chimpanzee searches and follows. We processed samples using water separation techniques to isolate undigested materials including seeds, leaf fragments, and fruit remnants. We categorized all identifiable parts by species, and recorded quantitative data recorded on seed counts and fiber content. Photographic documentation supported taxonomic classification.

## Data analyses and statistical Analyses

All analyses were conducted using R version 4.1.1 (R Core Team 2018) and can be found at the following link: https://github.com/ChimpanzeeResearch/bugoma_ecology.

### Dietary preferences

We assessed dietary preferences using chi-square tests for both species, item and food part preferences (we refer to parts for a specific part of the plant eaten; we additionally refer to item when it is a specific item in the diet, e.g. meat – note that a part thus constitutes an item and is listed as such in some of the figures). We used analysis of variance (ANOVA) to assess seasonal effects on fruit consumption comparing Ugandan typically defined seasons (two wet seasons: March-May and September-November; two dry seasons: December-February and June-August).

### Ripe fruit index (RFI)

We calculated the RFI as the proportion of ripe fruit consumption relative to all food items consumed. We calculated rolling averages of the RFI at multiple temporal scales (2, 7, 30, 49, and 90 days) to examine dietary patterns across different timescales.

### Species Diversity and Species Richness

To allow comparison with previous analyses of other Ugandan chimpanzee communities’ diets (Gruber et al., 2012), we quantified dietary diversity using the Shannon–Wiener index (*H* = − ∑_*i*_ *p*_*i*_ log *p*_*i*_), where *p*_*i*_ represents the proportion of species *i* in the overall diet. Higher *H* values indicate greater diversity. We also calculated the Hill’s equality index, *J*′, a standardized version of the Shannon–Wiener index, where *J*^′^ = *H*/ log *S*, with *S* being the total number of species in the diet during the sampling period (Hill, 1973). *J*′ ranges from 0 and 1, with 1 indicating uniform feeding across species.

For temporal analyses, we also calculated Species Diversity, using rolling windows as represented of Hill’s equality index values at multiple temporal scales (2, 7, 30, 49, and 90 days) to match the scales used for RFI analyses. In addition, for the Methodological part, we calculated Species Richness as the total unique food species identified through both observational and fecal sampling methods.

### Temporal Patterns and Weather Conditions

We examined weather effects including rainfall, humidity, and temperature variables on feeding behavior through Pearson’s correlation analyses at multiple temporal scales. We calculated rolling averages (2, 7, 30, 49, and 90 days) to account for cumulative effects. We examined correlations between the rolling weather averages and both RFI and Species Diversity at matching timescales, particularly in how the strength and direction of these relationships varied with the temporal window of analysis.

### Multiscale Predictive Models

To examine the relationships between dietary patterns and environmental variables across different temporal scales, we constructed linear regression models at five temporal windows (2, 7, 30, 49, and 90 days) for both the RFI and Species Diversity models. Year was included as a random effect in all models to account for potential habituation effects and temporal dependencies in the data. For both model types, we compared model performance across timescales using Akaike Information Criterion (AIC) and evaluated the consistency of predictor effects across scales. We constructed models with predictors at each timescale using rolling average values.

### Species Diversity Models

These models predicted dietary diversity using rainfall and temperature as continuous predictors, with year as a random effect. Both weather variables were standardized (z– scored) to allow comparison of effect sizes across predictors. We constructed models for each timescale using the rolling average values of rainfall and temperature.

### RFI Models

These models predicted 2-day RFI using Species Diversity index, rainfall, and temperature as predictors, with year as a random effect. All continuous predictors were standardized. We used Beta regression with a logit link to account for the bounded nature of the RFI (values between 0 and 1), using the glmmTMB package (Brooks et al. 2017).

### Comparison Between Observational and Fecal Sampling Methods

We analyzed differences in species detection and dietary composition between observational and fecal sampling methods using Fisher’s Exact Tests and Chi-square tests with Yates’ continuity correction. We calculated both diversity indices (Shannon-Wiener and Hill’s equality) separately for each sampling method to assess methodological differences in species detection capabilities.

For meat consumption, we compared detection rates between methods using both Fisher’s Exact Test and Chi-square analysis. We conducted two comparisons: (1) a full dataset comparison including all 964 observation days and 75 dung samples collected across 55 days, and (2) a matched subset analysis comparing only dates where fecal samples were collected with observational data from the previous day, allowing direct comparison of the same feeding events detected by different methods.

### Ugandan population comparison

We describe and compare dietary composition and diversity metrics from the Mwera South community with published data from other Ugandan chimpanzee communities (Kanyawara, Ngogo, and Sonso). The Sonso community (01°43’N, 31°32’E) is found in Budongo Forest Reserve, a 482 km² expanse of medium-altitude semi-deciduous forest, approximately 75km from the Mwera South community. The Kanyawara community (00°33’N, 30°21’E) is situated in Kibale National Park, a 795-km^2^ area mostly covered by moist evergreen or semi-deciduous forest, approximately 110km from the Mwera South community. Data from these communities were taken from Gruber et al. (2012).

We compared J’ across sites, noting that durations of the available data varied: Kanyawara (12 months), Ngogo (12 months), Sonso (15 months), and Mwera South (38 months), as did habituation of and contact time across the communities (research started with Kanyawara, Ngogo and Sonso in the 1990s, and with Mwera South in 2015). We compared diet composition across six major food categories (ripe fruits, unripe fruits, flowers, leaves, terrestrial herbaceous vegetation, and other foods) using percentage of feeding time for other sites and percentage of feeding days for Mwera South. We also identified shared tree species, including fig species across communities.

## Results

### Environmental variations

#### Temporal Patterns and Weather Conditions

Weather conditions in Bugoma Forest followed distinct seasonal patterns throughout the study period (Figure 1). Daily rainfall patterns aligned with the typically defined seasons in Uganda (see methods), with peaks during both wet seasons. Temperature and humidity displayed their own temporal variations, with humidity showing stronger seasonal fluctuations than temperature.

**Figure 1.**
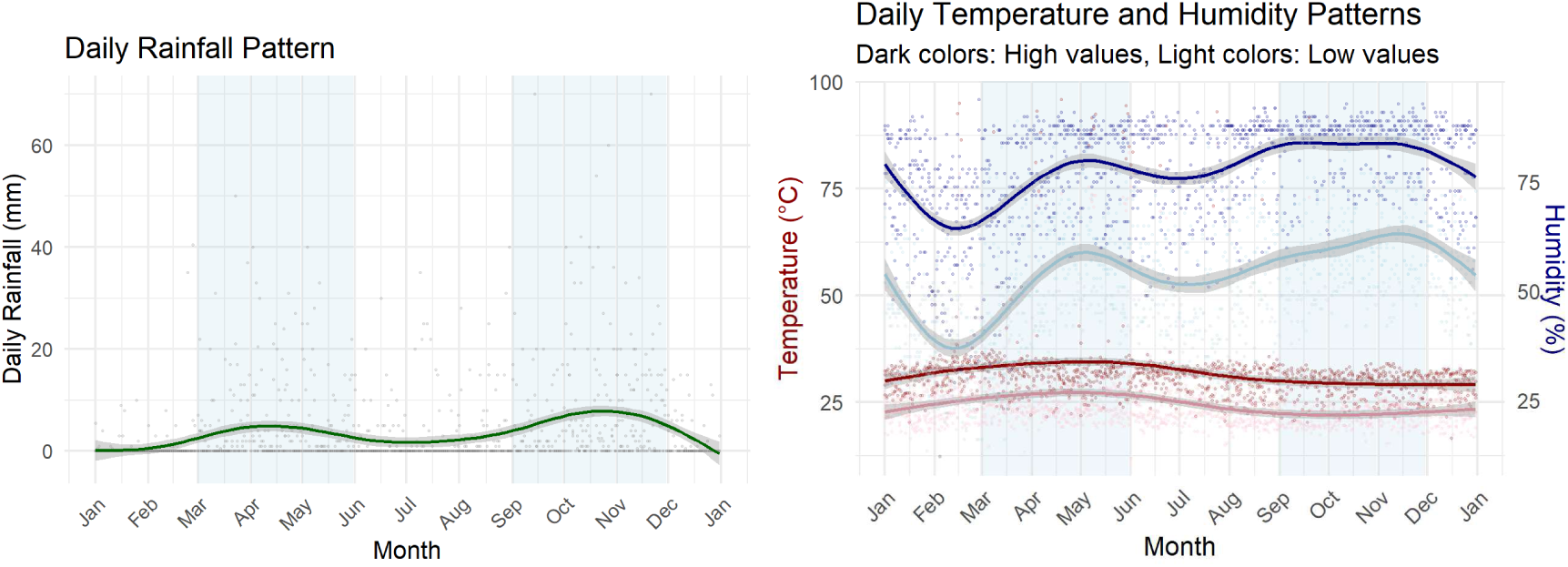
Meteorological variation over four years at the Mwera field station, in Bugoma Forest. The typically defined seasons in Uganda are depicted with blue shading for rainy seasons and white for dry seasons. Left: Daily rainfall patterns. Right: Daily temperature (in Celsius) and Humidity percentage patterns.

A weak positive correlation was found between rainfall and humidity (r = 0.23, 95% CI [0.19, 0.27], p < 0.001). Temperature showed no correlation with either rainfall (r = –0.03, p = 0.220) or humidity (r = –0.03, p = 0.103), suggesting relative independence of these parameters in Bugoma CFR.

#### Dietary diversity and seasonal variation in consumed foods

The Mwera South chimpanzee community exhibited clear dietary preferences across food items (Figure 2). The community consumed at least 66 different species in their diet (Figure S1). We further quantified this diversity using the Shannon-Weiner diversity index, which yielded a value of H = 3.09, indicating substantial dietary diversity. The Hill’s equality index, or the normalized Shannon-Weiner diversity index, was J’ = 0.74 (38 months). Fig trees were consistently among the top species consumed, as seen in other Ugandan communities (Figure S1, Table ST3). The Mwera South community showed significant preferences across food items (χ² = 15552, df = 17, p < 0.001), with distinct seasonal variations in the pattern of consumption (Figure 2). We found a strong negative correlation between (total) fruit consumption and leaf consumption (r = –0.75, 95% CI [– 0.77, –0.73], p < 0.001). This negative relationship was also evident, although slightly weaker, when examining ripe fruit consumption versus leaf consumption (r = –0.64, 95% CI [-0.66, –0.62], p < 0.001).

**Figure 2.**
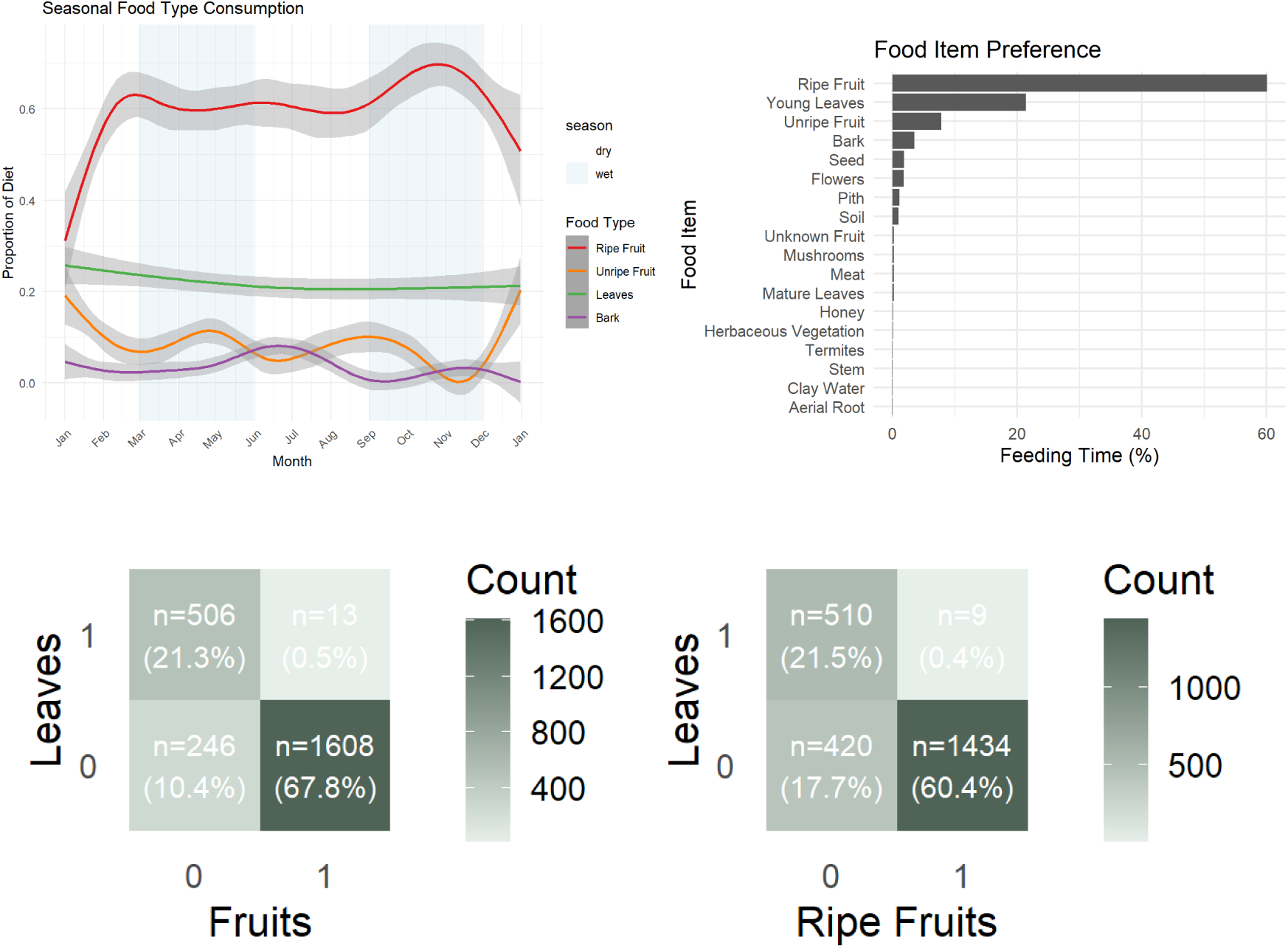
**Top**: The proportion of the preferred food items eaten throughout the year and the time spent feeding on a specific food item by the Mwera South chimpanzees. Seasons are depicted with blue shading for rainy seasons and white for dry seasons. Feeding time represents the frequency of days on which the chimpanzees consumed a particular food item. **Bottom:** The relationship between leaves and (total) fruit consumed (on left) and with ripe fruits only (on right).

### Timescale analyses on the relationships between ripe fruit consumption, species diversity, and environmental variables

#### Correlations between ripe fruit consumption and species diversity

We found a negative correlation between the Ripe Fruit Index and Species Diversity at all timescales (here: 7 days, r = –0.286, 95% CI [-0.46, –0.19], p < 0.001, Figure 3; additional timescales in Supplementary Figure S2).

**Figure 3.**
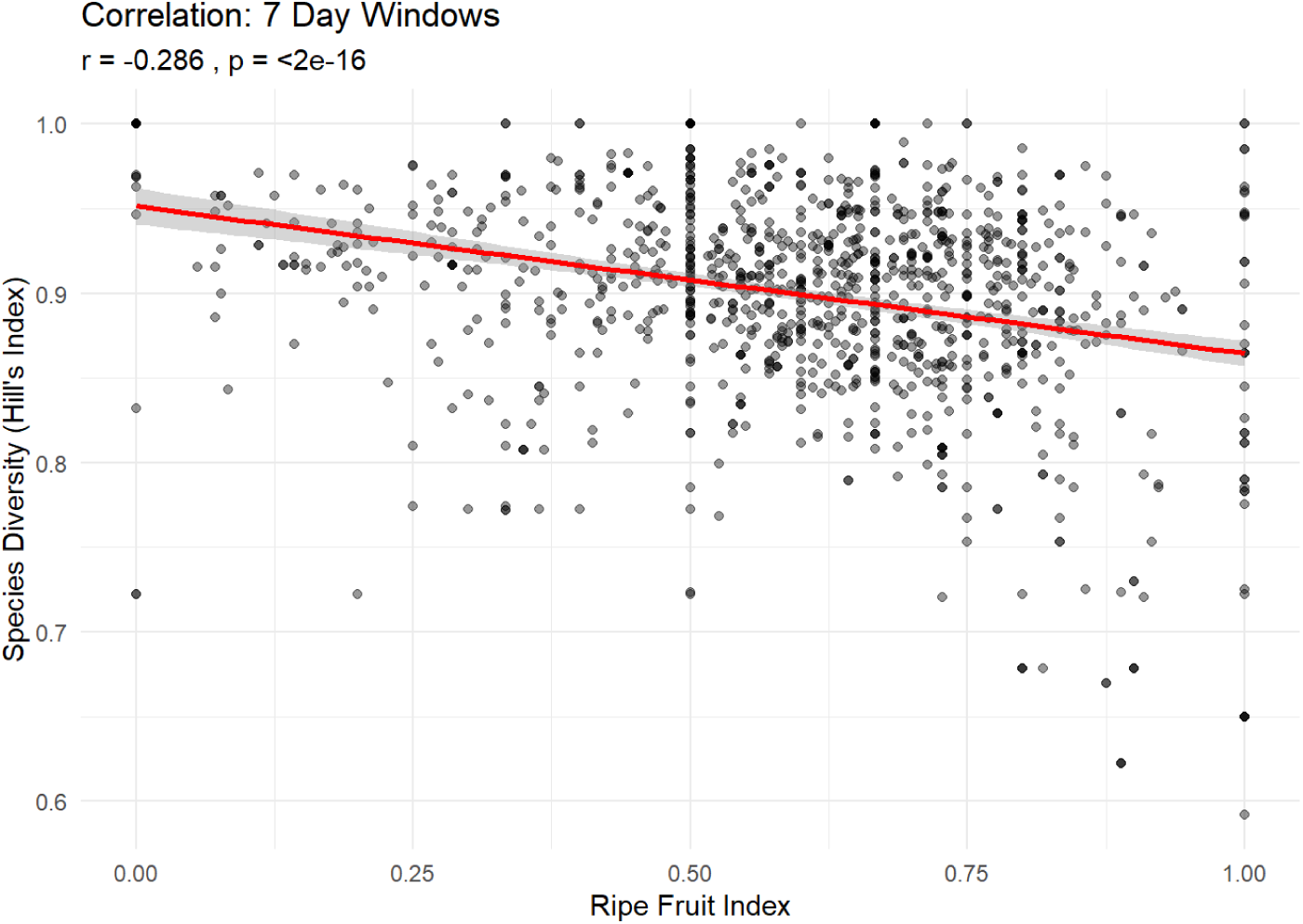
Correlation between the species diversity calculated by the Hill’s Index and the Ripe Fruit Index calculated with rolling averages of 7 days.

#### Correlations between Ripe Fruit Index and Species Diversity with Rainfall and Temperature

We correlated our dietary indices at various timescales to assess whether rolling averages of our indices correlated more strongly with average temperature or rainfall. While doing so inflates re-uses of the same values over time, we observed similar effects across the various time periods considered. Across timescales, the correlations between Ripe Fruit Index and Species Diversity with rainfall and temperature are quite weak (Supplementary Figure S3), with a (weakly) positive relationship between Ripe Fruit Index and rainfall (more ripe fruits correlate with greater rainfall) but a (weakly) negative relationship between Species Diversity and rainfall (less diversity with greater rainfall; Table 1). Temperature correlated negatively with both indices (Table 1).

**Table 1.** Correlations between rolling averages of Ripe Fruit Index (RFI) and Species Diversity, with rainfall and temperature across timescales.

| Time scale | RFI vs Rainfall |  | Species Diversity vs Rainfall |  | RFI vs Temperature |  | Species Diversity vs Temperature |  |
| --- | --- | --- | --- | --- | --- | --- | --- | --- |
|  | r | p | r | p | r | p | r | p |
| **2 Day** | 0.057 | 0.0872 | -0.043 | 0.194 | -0.129 | 0.0001 | 0 | 0.994 |
| **7 Day** | 0.127 | <0.001 | -0.033 | 0.272 | -0.112 | 0.0002 | -0.053 | 0.074 |
| **30 Day** | 0.225 | <0.001 | -0.133 | <0.001 | -0.108 | 0.0003 | -0.206 | <0.001 |
| **49 Day** | 0.297 | <0.001 | -0.158 | <0.001 | -0.159 | <0.001 | -0.208 | <0.001 |
| **90 Day** | 0.306 | <0.001 | -0.156 | <0.001 | -0.25 | <0.001 | -0.068 | <0.001 |

#### Multiscale predictive models

##### Species Diversity Index Models

Both rainfall and temperature were consistently negative across all timescales. The effects increased in strength from shorter to longer timescales (Table 2, see Supplementary Table ST1 for all timescales), with the model with the best fit provided by the longest timescale (Supplementary Figure S4; values at scale = 7 days are given both as reference, and because they provide the best fit for Ripe Fruit Index).

**Table 2.** Multiscale comparison of models’ prediction for Species Diversity index at 7 and 90 days using the rolling averages values of rainfall and temperature.

| *Timescale* | *Predictor* | *Estimate* | *SE* | *t* | *p* |
| --- | --- | --- | --- | --- | --- |
| 7 | Intercept | 0.897 | 0.008 | 115.188 | <0.001 |
| 7 | Rainfall | -0.005 | 0.002 | -2.488 | 0.0130 |
| 7 | Temperature | -0.008 | 0.002 | -3.871 | 0.0001 |
| 90 | Intercept | 0.813 | 0.021 | 39.261 | <0.001 |
| 90 | Rainfall | -0.013 | 0.001 | -13.497 | <0.001 |
| 90 | Temperature | -0.028 | 0.001 | -21.656 | <0.001 |

##### Ripe Fruit Index Models

These models predict 2-day Ripe Fruit Index using Species Diversity, rainfall and temperature. The model with the best fit was for a 7-day timescale (Table 3). The Species Diversity effect started out strongly negative at short timescales but completely disappeared by 30 days and beyond. The rainfall and temperature values showed more variation, with either negative or positive directions that became inverted as the timescale increased (see Supplementary Table ST2).

**Table 3.**
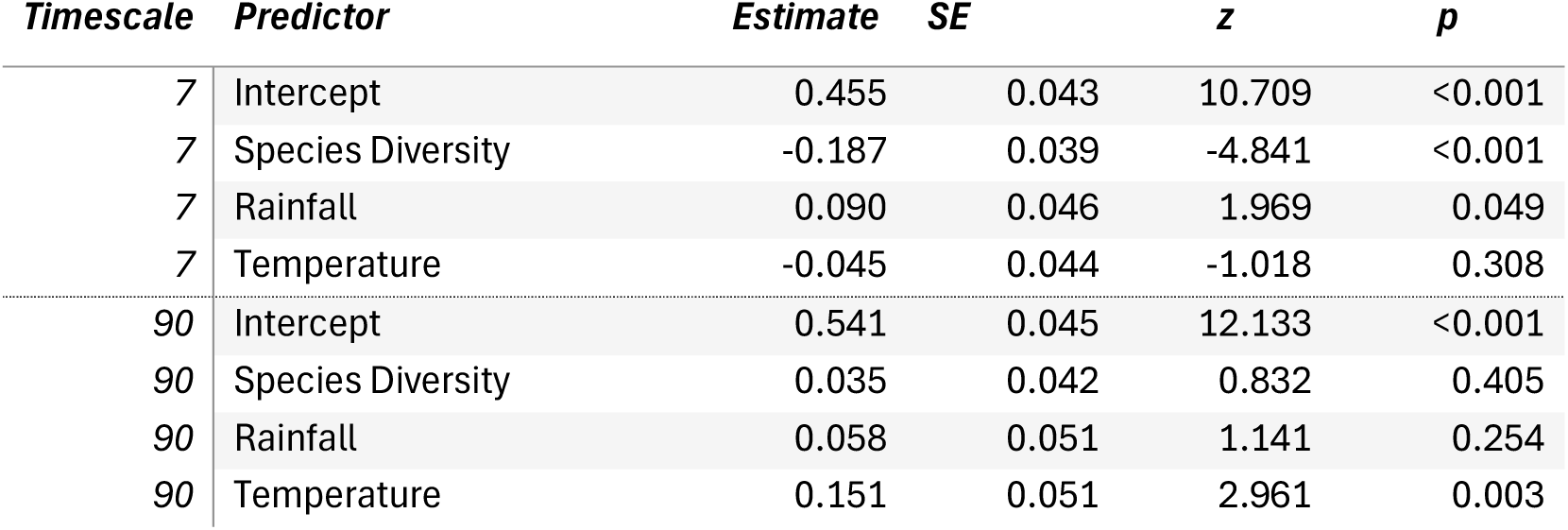
Multiscale comparison of model prediction for 2-day Ripe Fruit Index using 7– and 90– day rolling averages of rainfall, temperature, and Species Diversity.

| *Timescale* | *Predictor* | *Estimate* | *SE* | *z* | *p* |
| --- | --- | --- | --- | --- | --- |
| 7 | Intercept | 0.455 | 0.043 | 10.709 | <0.001 |
| 7 | Species Diversity | -0.187 | 0.039 | -4.841 | <0.001 |
| 7 | Rainfall | 0.090 | 0.046 | 1.969 | 0.049 |
| 7 | Temperature | -0.045 | 0.044 | -1.018 | 0.308 |
| 90 | Intercept | 0.541 | 0.045 | 12.133 | <0.001 |
| 90 | Species Diversity | 0.035 | 0.042 | 0.832 | 0.405 |
| 90 | Rainfall | 0.058 | 0.051 | 1.141 | 0.254 |
| 90 | Temperature | 0.151 | 0.051 | 2.961 | 0.003 |

### Methodological aspects

#### Comparison of dietary diversity using two sampling methods

The dung samples (N=75) contained 276 food items (∼3.7 food items per sample), compared to 2373 food items records across 964 observation days (∼2.5 food items per day). Direct observations and dung analysis revealed different patterns in both Species composition and Food item proportions (Table 4, Figure 4). The two methods showed varying success in detecting different dietary components. Species identification was substantially underestimated in direct observations (dung analysis identified 43 species, as compared to 23 species through direct observation; Table 4). Nevertheless, we found similar Hill’s equality indices (J’) between methods (0.83 and 0.86) indicating that, despite differences in Species Richness, both approaches captured comparable patterns in the relative evenness of food species consumption.

**Figure 4.**
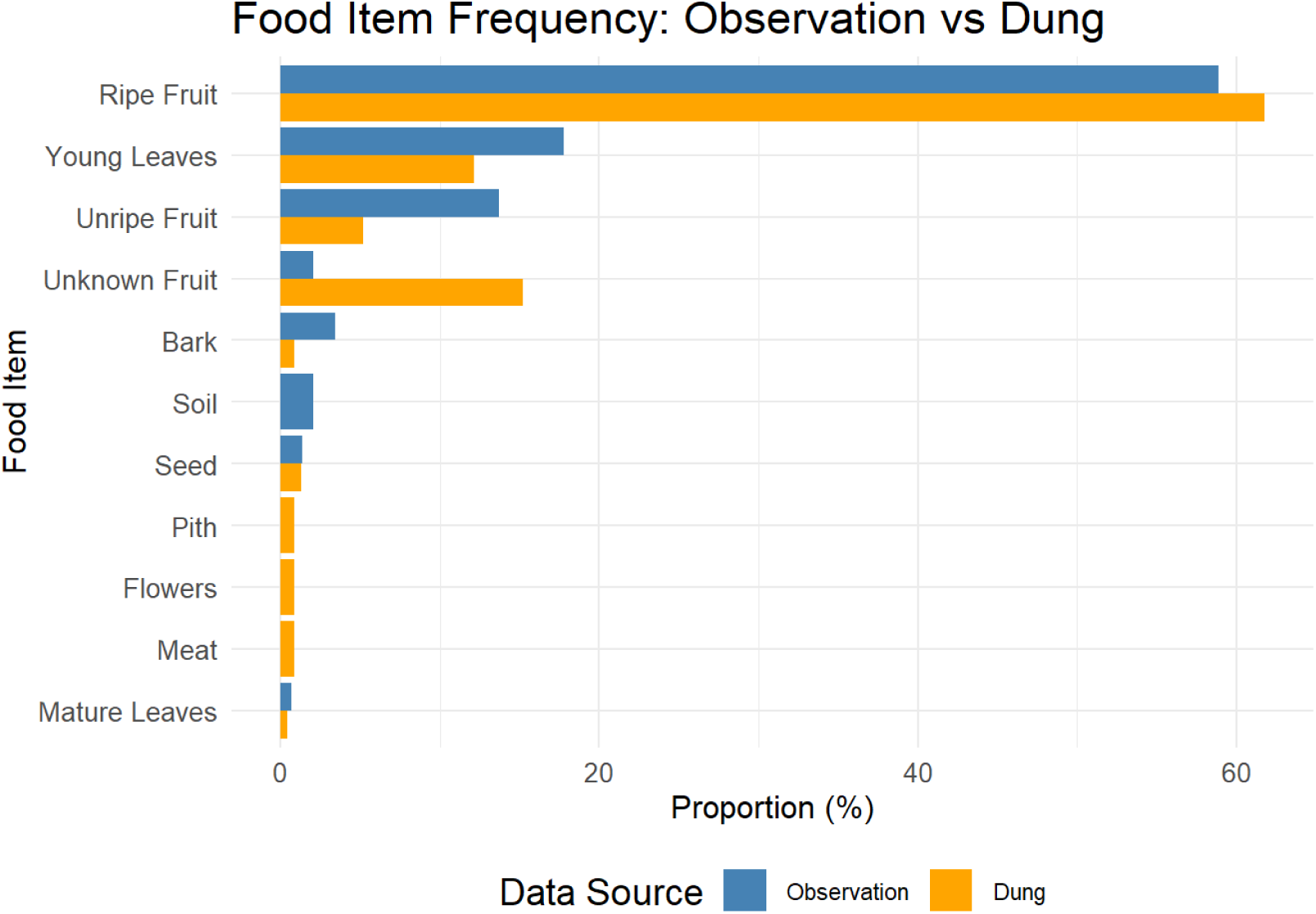
The proportion of food items consumed based on two different methods of data collection. The species identified in direct observations are in blue and those identified in dung sampling are in yellow.

**Table 4.**
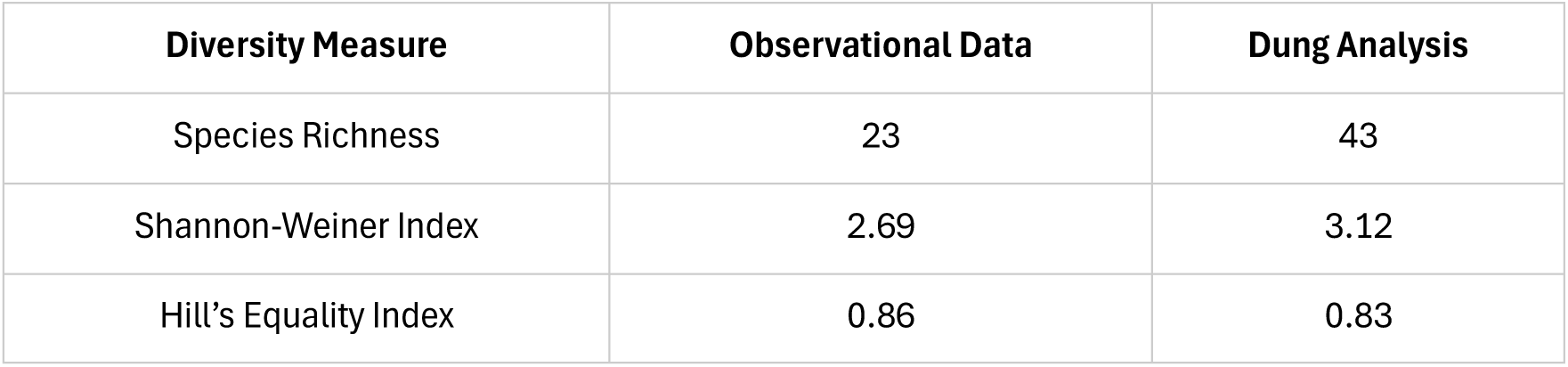
Comparison of Dietary Diversity Indices between Sampling Methods.

| Diversity Measure | Observational Data | Dung Analysis |
| --- | --- | --- |
| Species Richness | 23 | 43 |
| Shannon-Weiner Index | 2.69 | 3.12 |
| Hill's Equality Index | 0.86 | 0.83 |

#### Meat consumption

Meat consumption was also detected at different rates (Table 5). In our subset of matched samples, we detected meat consumption through dung sampling on three days, with no meat consumption detected in the observational data subset. In the full datasets for both sampling methods, direct observations recorded meat consumption on five occasions out of 2373 feeding observations (0.21%), while dung analysis identified meat remains in four out of 276 food item records in 75 samples (1.45%). Although the sample is small, direct observation appears to underestimate the frequency of meat consumption (Fisher’s Exact Test: odds ratio = 0.14, 95% CI [0.03, 0.73], p = 0.009; Chi– square analysis: χ² = 7.84, df = 1, p = 0.005). However, direct observations provided additional important qualitative data, such as the identification of Black-and-White colobus (*Colobus guereza*) as a prey species, which was not possible through dung analysis alone.

**Table 5.** Comparison of meat detection rates across food item records.

|  | Meat | No meat | Proportion |
| --- | --- | --- | --- |
| Direct observation | 5 | 2368 | 0.21% |
| Dung sampling | 4 | 272 | 1.45% |

Values reported are individual food item records.

### Comparison with other Ugandan field sites

The J’ value for the Mwera South community (0.74, over 38 months) is comparable to the values reported for other chimpanzee communities in Uganda. For example, Kanyawara chimpanzees had a J’ value of 0.70 (over 12 months), and Sonso chimpanzees had a J’ value of 0.69 (over 15 months), whereas Ngogo showed a slightly lower value at 0.58 (over 12 months).

A large number of tree species present in the diet were shared across communities, particularly fig species (e.g.: *Ficus mucuso*, *F. natalensis* and *F. sur/capensis*; for a full comparison see Supplemental Table ST3). All chimpanzee communities showed a preference for particular food items (χ² = 15552, df = 17, p < 0.001), with a strong preference for ripe fruits and leaves (Table 6).

**Table 6.** Diet composition across four chimpanzee study sites in Uganda shown as percentage of feeding time. Diet composition is broken down into six major food items: ripe fruits, unripe fruits, flowers, leaves, terrestrial herbaceous vegetation (THV), and others. This table adapted from Gruber et al. 2012, which summarized 15 months of data for Sonso (extracted from Newton-Fisher, 1999), and 12 months of data for Kanyawara and Ngogo (extracted from Potts et al., 2011). Bugoma data represents observational feeding records collected through focal animal sampling (April 2021 to May 2024). Note that in Kanyawara, Ngogo, and Sonso percentages represent proportion of feeding time, whereas in Bugoma percentages represent proportion of feeding days.

| Food item | Kanyawara | Ngogo | Sonso | Bugoma |
| --- | --- | --- | --- | --- |
| Ripe fruits | 64.6 | 80.5 | 54.6 | 60.4 |
| Unripe fruits | 2.0 | 11.0 | 9.9 | 7.9 |
| Flowers | Included in Others | Included in Others | 8.8 | 1.8 |
| Leaves | 11.4 | 3.5 | 19.7 | 21.6 |
| THV | 17.4 | 1.1 | 3.2 | 1.3 |
| Others | 4.6 | 3.9 | 3.8 | 7.0 |
| Total | 100 | 100 | 100 | 100 |

## Discussion

Primates, particularly great apes, exhibit complex feeding strategies that reflect both ecological constraints and behavioral adaptations to their environment. This relationship is crucial for large-bodied primates, whose substantial energetic requirements and relatively long life histories necessitate efficient foraging strategies (Aiello & Wheeler, 1995). Our study provides the first comprehensive characterization of feeding ecology in the Mwera South chimpanzee community of the Bugoma Cental Forest Reserve, Uganda, revealing patterns of feeding ecology that both align with and diverge from those described in other Ugandan populations. As predicted in H4, the community showed clear dietary preferences, with a small number of species constituting the majority of their diet (Figure 2 and Supplementary Table 3). The strong preferences we documented for specific food items, in particular young leaves of *Celtis midbraedi* and the ripe fruits of several species of fig, demonstrates remarkable dietary selectivity, but also reflects unremarkably observations in other Ugandan communities. We review our findings in terms of three major sources of potential influence: environmental aspects and timescales, methodological challenges, and cultural aspects.

### Environmental aspects and timescales

As predicted in H2, we found clear correlations connected to the presence of fruits across our analyses. These relationships suggested a dietary switching pattern in which increased fruit consumption was associated with decreased leaf consumption, and vice versa. The stronger negative correlation between all fruit consumption and leaves, as compared to ripe fruit consumption alone indicates that when the Mwera South chimpanzees were transitioning away from leaf consumption, they supplemented their diet with unripe fruits, rather than relying solely on ripe fruits. When ripe fruits were available, chimpanzees fed on them almost exclusively, suggesting strategic dietary choices that optimized energy intake while minimizing foraging effort, as seen in other populations (Watts et al., 2012). These findings are also supported by our multiscale models. Regarding Species Diversity, we found consistent effects of rainfall and temperature across all timescales from 7-day onward: both were consistently negative, meaning that more rainfall and higher temperatures were associated with lower dietary diversity. In the Ripe Fruit Index models, Species Diversity had the most consistent effect at short timescales, being strongly negative (with the best fit found at a 7-day timescale), and illustrating here again that the more chimpanzees fed on ripe fruits, the less diversity they had in their diet. However, the fact that Species Diversity effects disappeared by 30 days suggest that the relationship between Ripe Fruit feeding and Species Diversity should be understood as an ad-hoc consequence (‘there is one ripe fruit spot, let’s focus on it’) rather than as a feeding strategy (e.g. ‘if low diversity over the last month, focus on ripe fruits in the future’). Overall, our results suggest that several aspects of dietary diversity, whether at the level of Species or ripe food proportions have to be collected; and that their relationship with each other, as with other ecological indicators such as temperature or rainfall, is not necessarily trivial, nor equivalent across timescales.

### Methodological challenges

Our multi-method approach to studying feeding ecology tackled the issue of the influence of methods on results. Our results strongly supported H3, with dung analyses revealing higher species richness than observational data alone. Our results underline the value of dung analyses, particularly in newly habituated populations where direct observations may miss specific feeding events. Of particular note, our dung analyses suggested higher rates of meat consumption than recorded through observational data. The complementary insights gained from both methods revealed important aspects of dietary composition that would have been missed with a single approach. Dung analyses offered a more complete picture of actual consumption patterns and appear particularly important to capture feeding events outside the observation window (Phillips & McGrew, 2013), especially where and when habituation to direct observation is ongoing. At the same time, observational data provided crucial context about hunting behavior and prey species. The apparent difference in species detection between methods may also reflect the capacity of observers to detect species through direct observation, particularly at relatively new field sites. Our observational data necessarily grouped certain plant types (particularly climbers) into broad categories, due to the challenges of field identification of living plants, whereas dung analyses allowed more precise differentiation through seed morphology.

Finally, given the similar Hill’s equality indices between methods (0.83 and 0.86), despite differences in species richness, both approaches captured comparable patterns in the relative evenness of food species consumption. As a result, the general pattern of dietary intake remains consistent across methodologies, and while future use of additional methods such as eDNA could allow greater granularity in species specificity, it is unlikely to substantially change the overall pattern.

### Cultural aspects

Mwera South chimpanzees shared dietary preferences with both Budongo and Kibale communities, particularly their heavy reliance on figs, supporting H5 about regional dietary overlap. In particular, the comparison of J’ values suggested that Mwera South community maintained a relatively high dietary diversity, similar to or slightly higher than other studied communities.

#### The Cultural Junction Hypothesis

The geographical location of Bugoma Forest between several well-studied chimpanzee populations in Uganda and the Democratic Republic of Congo provides a unique opportunity to examine difference and similarities between communities that may be considered cultural. Historically, Uganda was widely forested but, in recent history, forest cover has greatly reduced (Gruber, 2013; McLennan et al., 2020) and the remaining forest reserves and patches are fragmented, restricting movement between them. Migration between chimpanzee communities offers a route for the transmission of cultural behavior (Luncz & Boesch, 2014; Péter et al., 2022), but only where contiguous habitat allows natural migrations to occur. The fragmentation of forests across Africa impacts not only the cultural transmission of chimpanzee behavior, but also the genetic diversity of chimpanzee populations—and potentially their long-term survival (Kühl et al., 2017).

Bugoma chimpanzees exhibit a distinctive behavioral profile that includes traits from both Ugandan and Congolese populations, suggesting that this area may have served as a ‘cultural junction’ in the past. At the same time, our ability to assess this hypothesis is limited where we are unable to rule out alternative explanations (Laland & Janik, 2006; Whiten et al., 2001)—such as patterns of similarity and distinction in local ecology. Nevertheless, ecological differences can themselves provide the seeds for cultural differences, a growingly acknowledged position in the literature (Gruber et al., 2012; Kalan et al., 2020; Möbius et al., 2008; Schöning et al., 2008).

While a first description, our ecological data support the argument for cultural, as well as ecological, influences on the diets of Ugandan chimpanzee communities. We find striking similarities in general dietary patterns between the Mwera South community and neighboring populations—as evident in aspects such as a comparable Hill’s equality index, which do not need cultural explanations. Yet acknowledging that the dietary preferences and patterns of food availability in Bugoma share characteristics with both Budongo and Kibale communities also suggest that environmental influences are unlikely to fully explain behavioral differences between these populations, particularly with respect to their tool use behavior (Mannion et al., 2025).

In effect, our findings on diet connect with recent evidence for cultural differences in the use of tools for extractive foraging across these communities. Mwera South chimpanzees employ both stick tools and leaf-sponges for honey extraction (Mannion et al., 2025). Stick tool use happens in other Ugandan communities, including the nearby Bulindi, Kasongoire, and Kasokwa chimpanzees in the Budongo-Bugoma corridor (Oxley & Jovan, 2019), as well as in the more distant Kibale (Watts, 2008) and Kalinzu populations (Koops et al., 2015). While other chimpanzee communities habitually employ stick tool use in foraging, evidence within Bugoma has remained scarce. At the same time, these occasional observations stand in stark contrast to the Budongo populations, in which stick use has remained firmly absent in feeding context despite over 35 years of observation and a series of field experiments (Gruber, 2016; Gruber et al., 2009). Overall, these patterns of behavioral differences in non-tool assisted and tool-assisted foraging between the Ugandan chimpanzee populations, advocates for both cultural and ecological influences as drivers of diets. The combination of cultural traits in Bugoma chimpanzees, including their diet, and the unique combination of tool use techniques and ground nesting behaviors typically associated with Congolese communities (Hobaiter et al., 2024; Romani et al., 2023), suggests that Bugoma chimpanzees may have sat at a historic junction between connected forests that permitted cultural exchange between what are—today—fragmented Ugandan and Congolese populations.

## Conclusion

Our characterization of Bugoma chimpanzee feeding ecology provides a foundation for future research at this site, as well as important context for the wider tapestry of Eastern chimpanzee populations. The selective behavior of Bugoma chimpanzees raises intriguing questions about how such preferences develop and are maintained within populations, particularly as chimpanzees navigate increasingly human-modified landscapes in these areas, leading to potential cultural adaptations (Gruber et al., 2019; McCarthy et al., 2017; McLennan & Hockings, 2014). The evidence for the role of the Bugoma Forest as a possible cultural junction between characteristically Ugandan and Congolese behavioral repertoires emphasizes the importance of maintaining habitat connectivity for preserving both behavioral diversity and genetic exchange (McLennan et al., 2020; Plumptre et al., 2021). As anthropogenic pressures on East African forests increase, understanding the flexibility and constraints in chimpanzee feeding ecology becomes crucial for effective locally-specific conservation strategies (Wessling et al., 2025). This understanding should inform development plans that balance human needs with the preservation of these unique populations and their behavioral traditions. By documenting both the ecological similarities and behavioral differences between Ugandan chimpanzee communities, we contribute to a more nuanced understanding of how culture and ecology interact to shape primate behavior—knowledge that is essential for preserving both the populations and their unique behavioral heritage.

## AUTHOR CONTRIBUTIONS

Kelly Ray Mannion (KRM) and Thibaud Gruber (TG) conceived the ideas and led the writing of the manuscript. KRM collected all data at BPCP with assistance from BPCP field assistants. TG and Catherine Hobaiter (CH) have founded and co-direct research and conservation activities at the Bugoma Primate Conservation Project. All authors contributed critically to the drafts and gave final approval for publication.

## ACKNOWLEDGEMENTS

Kelly Ray Mannion and Thibaud Gruber were supported by a grant of the Swiss National Science Foundation (grant PCEFP1_186832 to Thibaud Gruber). We are greatly appreciative of the Bugoma Primate Conservation Project field assistants, staff and research coordinators for their help with fieldwork. The Bugoma Primate Conservation Project (BPCP) Field team was instrumental to the collection of the present data. At the time of project, the BPCP Field team was composed of: Adaku Amon, Ndora Michael, Kugonza Stephen, Owota Solomon, Arua Yolam, Abou Julius, Okongo John-Martin, Twinamasiko Emmanuel and the late Manyanja Gerald. We are grateful to Ugandan National Council for Science and Technology (UNCST), Uganda Wildlife Authority (UWA) and the National Forest Authority (NFA) for permissions to conduct this project in Bugoma Forest Reserve. We are grateful to Jane Goodall Institute Schweiz for their financial support of BPCP.

